# Paths to annihilation: Genetic and demographic consequences of range contraction patterns

**DOI:** 10.1101/2021.01.26.428313

**Authors:** Jordan Rogan, Mickey Ray Parker, Zachary B. Hancock, Alexis D. Earl, Erin K. Buchholtz, Kristina Chyn, Jason Martina, Lee A. Fitzgerald

**Author notes:** Mailing address*: Jordan Rogan, Department of Ecology and Conservation Biology, 534 John Kimbrough Blvd., TAMU 2258, Texas A&M University, College Station, TX 77843-2258. Statement of Authorship: JR, MRP, ADE, EKB, KC, JPM and LAF conceptualized the study. JR, MRP, ZBH, LAF wrote the manuscript. ZBH designed the models and performed the simulations. JR, MRP, ADE, EKB, KC, and JPM performed the literature review. All authors contributed to editing final drafts. Data accessibility statement: Code for all models and analyses can be found at https://github.com/hancockzb.

## Abstract

Species range contractions are important contributors to biological annihilation, yet typically do not receive the same attention as extinctions. Range contractions can lead to marked impacts on populations but are often only characterized by measurements of reduced extent. For effective conservation efforts, it is critical to recognize that not all range contractions are the same. We propose four distinct patterns of range contraction: *shrinkage*, *amputation*, *hollow*, and *fragmentation*. We tested their impact on populations of a generic generalist species using forward-time simulations. Results showed that all four patterns differentially reduced population abundance (declines of 60-80%) and significantly increased average relatedness, with differing patterns in nucleotide diversity (π) declines relative to the contraction pattern. The fragmentation pattern resulted in the strongest effects on post-contraction genetic diversity and structure. Defining and quantifying range contraction patterns and their consequences for the planet’s biodiversity provides necessary information to combat biological annihilation in the Anthropocene.

## Introduction

Widespread impoverishment of biodiversity, referred to as “biological annihilation” (Ceballos et al. 2017) is occurring across taxon groups and ecological scales. Yet focusing attention solely on extinctions underestimates the severity of the biodiversity crisis (Dirzo *et al*., 2014). Findings on range contraction for mammals by Ceballos *et al.* (2017) are alarming: nearly all of the 177 species they examined have lost 40% or more of their geographic range, with almost half losing more than 80%. In a separate analysis, Ceballos *et al.* (2020) found that, for 48 mammal and 29 bird species on the brink of extinction, there was an estimated reduction of 95% and 94% in their ranges since 1900, respectively. While the gravity of species extinctions is a compelling narrative within the biodiversity crisis, extinction accounts for only a small portion of overall biodiversity decline (Butchart *et al.,* 2010; Dirzo *et al.,* 2014).

Range contraction is typically described as the amount of range lost (Ceballos *et al.,* 2017, 2020). However, range contractions can take various patterns beyond the spatial extent of loss and each of these patterns can have different consequences for species’ populations. Distinguishing between range contraction patterns therefore has important implications for conservation. Different patterns may necessitate different strategies, such as, conserving disjunct populations, prioritizing conservation corridors, reintroduction planning, or restoring habitat within a species’ historic range. Effects of range contraction on population demography and genetics are understudied and may vary according to the contraction pattern.

Recent advances in simulation software (Kelleher *et al.,* 2018; Haller *et al.,* 2018; Haller & Messer, 2019) have expanded our ability to assess how range contractions impact populations. For example, by utilizing *tree sequence recording* (Haller *et al.,* 2019) in the forward-time simulator SLiM (Haller & Messer, 2019), it is possible to track the local genomic ancestry of every individual alive during the simulation. Combining this with the spatial locations of individuals, investigators can evaluate how range contraction patterns bias the distribution of ancestors backward in time.

Few, if any, studies have explored the potential consequences of distinct patterns of range contraction on demography and genetics of species’ populations. We hypothesize that distinct contraction patterns will produce different demographic and genetic consequences, and these consequences will not be uniform throughout post-contraction ranges. We investigated this hypothesis by simulating the demographic and genetic effects of four different range contraction patterns on a generalist species and evaluating the significance of these impacts for mitigating future species decline due to range contractions. In **Box 1**, we discuss the prevailing hypotheses of how ranges contract and how we used those, coupled with real-world examples (**Table 1**), to define the four patterns. The goal of our simulation models was to gain insights into the interplay between range contraction patterns and their consequences for genetic diversity and demography. In addition to tracking genetic diversity (π), we also mapped spatial ancestry. Finally, we examined how sampling individuals from different parts of the range can alter interpretation of the impacts of the contraction pattern. Our models allow us to make generalizable predictions about the magnitude and timing of effects of range contraction on genetic diversity and demography, and how the spatial distribution of ancestry and genetic diversity influence the interpretation of these effects. These insights can serve to inform the development of conservation interventions aimed at confronting the challenge of biological annihilation.

**Table 1.**
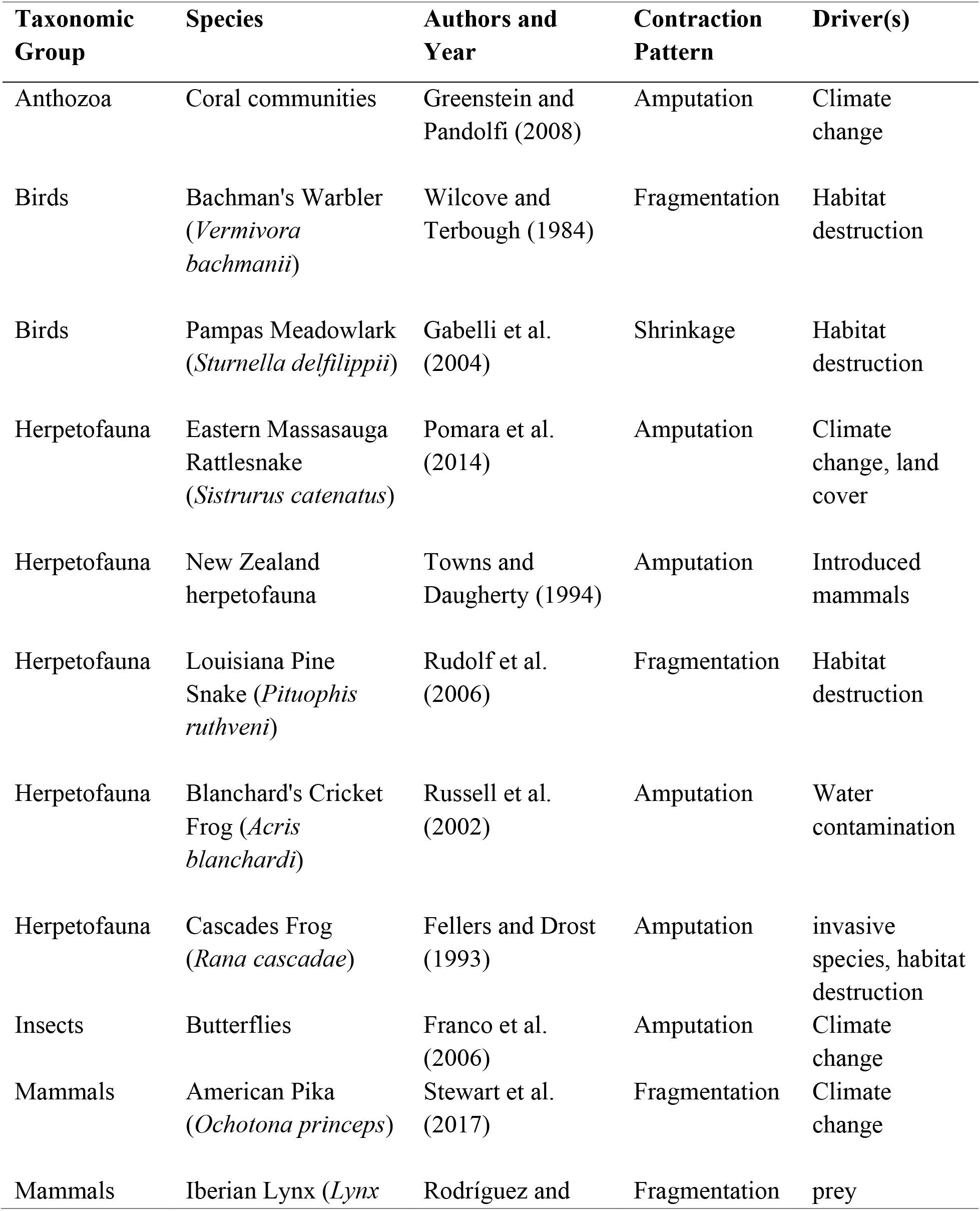
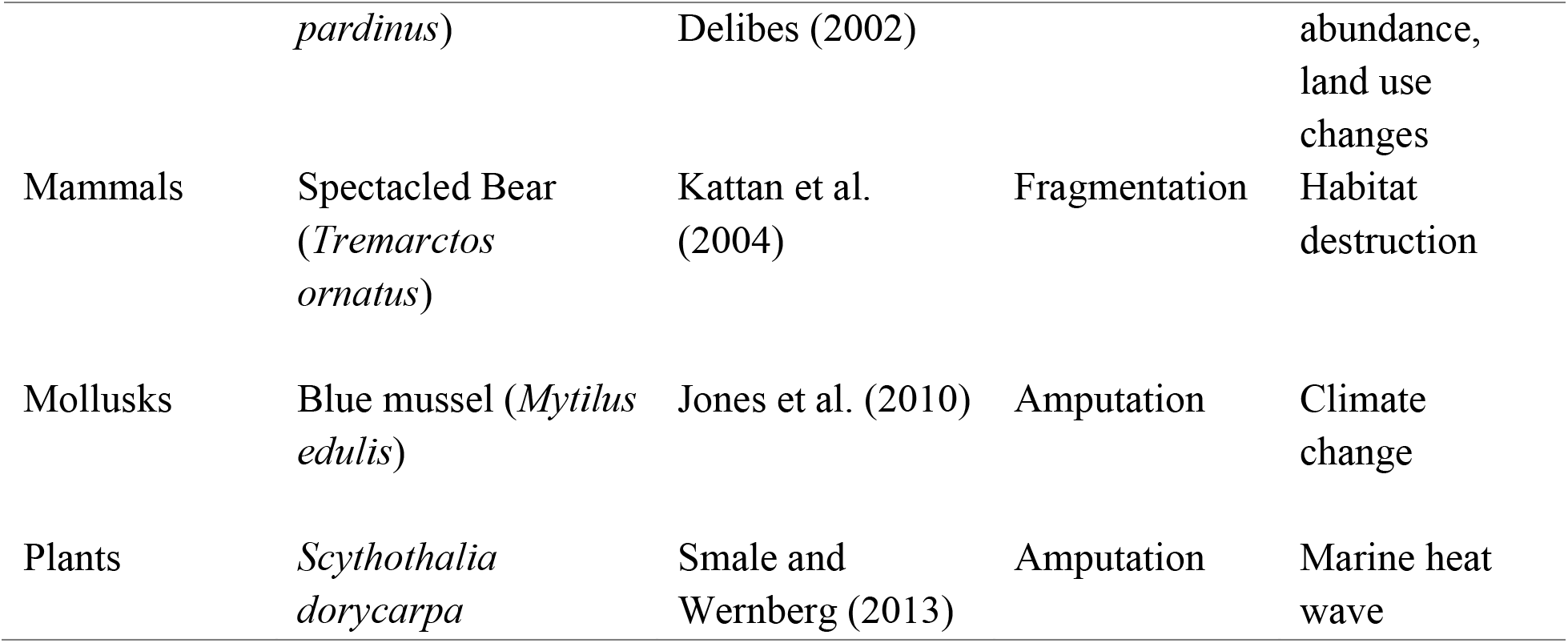
Examples of patterns of range contractions found in various taxa.

## Methods

### Population Model

We modelled impacts of four range contraction patterns (**Box 1**) on demographics and genetic diversity of a wide-ranging habitat generalist species using the forward-time simulation program SLiM v3.3 (Haller & Messer, 2019) (**Figure 1)**. Simulated individuals were diploid (2*n*), with haploid genome sizes of 1000 Mb and a uniform recombination rate of 10^−9^. Mutation was suppressed during the SLiM run and added later to decrease computation time (Kelleher *et al.,* 2018). Individuals were hermaphroditic, but self-incompatible. We modelled populations with overlapping generations in contiguous habitat with dimensionality (*x*, *y*). We included variation in the species’ distribution within its range by incorporating fitness values in 20×20 matrices, where unsuitable portions of the range have values of 0.1 (a 90% fitness reduction) and suitable have fitness values of 1.0 (i.e., no habitat-specific fitness cost).

**Figure 1.**
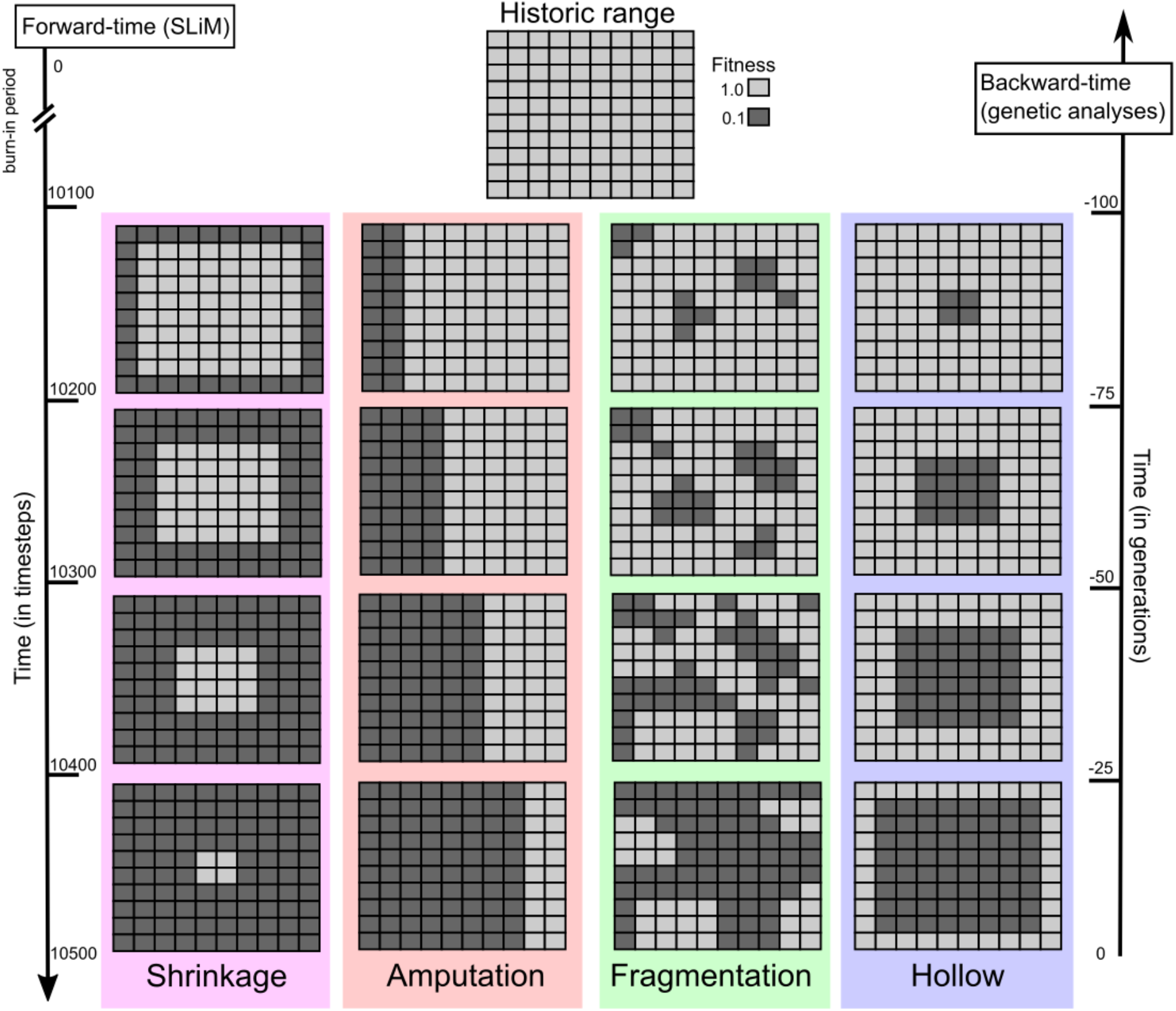
Simulation model showing both forward-in-time simulation and backward-in-time analysis directions. Size of the ranges have been scaled down to 10×10 instead of 20×20 for easier visualization of contraction patterns. While shown as a grid with distinct boundaries between cells, note that interpolation is used and there is a smooth gradient between them. Historic range has no cell-specific fitness reduction (though there are fitness consequences relative to distance from the edge).

At the beginning of each SliM run, 2000 individuals were randomly distributed across the range. Each timestep progressed through a series of events that included mate choice, offspring generation, and dispersal. Each of these processes were determined, to some degree, by a constant interaction distance, *σ* (= 0.5). We kept the interaction distance constant to reduce variation in results due to varying mate choice, dispersal, or spatial competition strength. For both mate choice and spatial competition, the interaction strengths were defined by a Gaussian distribution with a maximum of 1 / 2π*σ*^2^, where π is the mathematical constant, and standard deviation *σ*. The interaction strength had a maximum distance of 3*σ*, beyond which spatial competition and probability of mating are both effectively zero. Following Battey *et al*. (2020), the sum of all competitive interactions is *n_ij_* = *∑_j_ g(d_ij_)*, where *g* is a Gaussian density with mean 0 and standard deviation *σ* and *d* is distance from a given individual (again, max distance of 3*σ*). Mates were chosen randomly from within this maximum distance, and the number of offspring per mating pair was drawn from a Poisson distribution with *λ* = 1 / *L*, where *L* is the mean age at stationarity. Accounting for spatial competition and habitat quality (*h*), the absolute fitness of individual *i* (the probability of surviving to the next timestep) is calculated as:

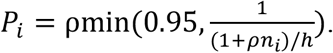

In the equation above, ρ = *F* / *K*(1 + *F*), where *F* is the mean fecundity and *K* is the local carrying-capacity. To avoid edge effects, we reduced fitness relative to distance from the edge (*W*):

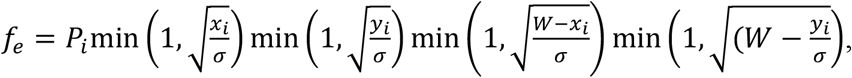

where *x_i_* and *y_i_* is the individual’s position in the *x* and *y* coordinates (Battey *et al.,* 2020; Bradburd & Ralph, 2019). Following mate choice and offspring generation, individuals dispersed according to a random normal distribution with mean zero and standard deviation *σ* in both *x* and *y* coordinates (following Battey *et al.,* 2020). All models had a burn-in period of 10,000 timesteps (~10*N*) before range contractions began.

We simulated each model of range contraction by altering the landscape of fitness values at four discrete intervals of 100 timesteps, each of which corresponds to ~25 (**Figure 1**). Each interval progressively decreased the total suitable habitat in a configuration according to the contraction pattern. To simulate a range shrinking as predicted by the demographic hypothesis, the *shrinkage* model initially reduced habitat suitability in the range’s periphery and continued reducing suitability towards the core at each interval. To simulate an extinction factor spreading across the landscape from its point of initial impact, as described by the contagion hypothesis, the *amputation* model initially reduced suitability in the periphery of the range, and at each interval this reduction expanded towards the periphery opposite the initial impact. The *hollow* model, also derived from the contagion hypothesis, initially reduced suitability in the core of the range, with suitability decreasing towards the periphery at each interval. The *fragmentation* model initially reduced suitability in several places in the range, with reductions in suitability expanding to create disjunct fragments. (See **Table S1** in Supporting Information for the total percent of range loss for each type of contraction).

During the run, SLiM tracked the local ancestry of each recombination breakpoint interval for all individuals via tree sequence recording (Haller *et al.,* 2019). In addition, we utilized SLiM’s ability to track the full pedigree of all individuals, which allowed us to estimate an average of Wright’s coefficient of relatedness (Wright, 1922). We did so by randomly sampling 50 individuals each generation, estimating their pedigree relatedness, and then estimating average sampled relatedness as: *F_r_* = (*r* – *n*) / *n* where *r* is the sum of all values in the relatedness matrix and *n* is the sample size.

The outputs of SLiM were the aforementioned tree sequences which store the complete population pedigree with each individual tagged in each timestep with their total number of offspring, age, fitness, and spatial location. Specific SLiM recipes for each contraction model are available at https://github.com/hancockzb.

### Genetic analysis

We analyzed genetic data from the SLiM model to determine how nucleotide diversity (π) and spatial ancestry were impacted by patterns of range contraction and how spatial sampling schemes impacted our interpretation. Nucleotide diversity is a common metric used in conservation genetics including inbreeding and population size estimation (e.g., Carvalho *et al.,* 2019, Hoelzel *et al.,* 1993; Roelke *et al.,* 1993), After each SLiM run, we analyzed the tree sequences produced using the python packages *tskit* and *pyslim* (Kelleher *et al.,* 2018), and *msprime* (Kelleher *et al.,* 2016). Trees were first checked for coalescence and, if multiple roots were found, we used the “recapitation” method in *pyslim* to simulate coalescence (Haller & Messer, 2019). We then overlaid neutral mutations (*μ* = 10^−8^) onto the trees, and performed all further analyses on the resulting tree sequences.

For each model of range contraction, we calculated π for 100 randomly selected individuals across four time points: 1) before the contraction began; 2) 25 generations after the contraction; 3) 50 generations after; and 4) 100 generations after. Additionally, we scaled π across time points relative to the pre-contraction mean, which enabled visualization of clusters with high or low diversity spatially. To determine if π was significantly less post-contraction, we performed a pairwise Wilcoxon test in the R platform (R Core Team, 2019). We used nonparametric tests throughout due to the data violating the assumptions of normality (Shapiro-Wilkes test, *p* < 4.1e-13).

To determine how spatial sampling schemes impacted our interpretation of the consequences of range contractions, we sampled 50 individuals each from 5–6 groups alive in the final generation from specific locations in the remaining range (“topleft,” “topright,” bottomleft,” “bottomright,” “center” for *shrinkage* and “top,” “uppermiddle,” “lowermiddle,” “lower” for *amputation*) and compared them to random samples of 50 individuals from the population prior to the contraction (“ancient”). The *hollow* and *fragmentation* scenarios had 5 groups (“topleft,” “topright,” bottomleft,” “bottomright,” “ancient”) because there was no “center” group. We computed pairwise nucleotide divergence (π12) between each group, as well as π within groups. We compared these results with a branch-length analogue statistic computed by *tskit* that does not rely on mutations, but instead computes the average distance (in number of timesteps) up the tree sequence between two randomly selected chromosomes within or between the specified group(s). In addition, we calculated pairwise *F*_ST_ for each group as

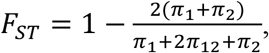

where π_1_ and π_2_ is the diversity within group 1 and 2, respectively.

To evaluate how patterns of spatial ancestry were impacted by range contraction, we randomly sampled 5 individuals alive at 25 generations (or 100 timesteps) after the end of the contraction and traced their ancestry back to the beginning of the contraction. This was performed in between recombination breakpoints by traversing up the tree to the ancestor(s) living in the given generation of interest. We tracked the proportion of the genome inherited from any single ancestor and recorded said ancestor’s spatial position. To determine the transition between the scattering (spatially autocorrelated ancestry) and collecting (random mating) phases (Wakeley, 1999), we compared the spread of spatial ancestry of the sampled individuals 25 generations before the contraction to a “neutral” model in which no contraction had occurred. This analysis allowed us to assess the relationship between the true dispersal distance (*σ*) and the effective dispersal distance (*σ*_e_). This analysis was performed using modified Python packages from Bradburd & Ralph (2019: https://github.com/gbradburd/spgr).

### Demographic analysis

We tracked the demographic parameters of age structure, mean number of offspring, and relative fitness throughout the SLiM simulation. Age was measured as the number of timesteps an individual survived through. We tested for a statistically significant relationship between each parameter and generation time via linear regression. We visualized areas of high and low relative fitness spatially by plotting individuals across four sampled time periods: 0, 25, 50, and 100 generations, where generation 0 is the population 25 generations before the contraction began and the 100th generation is 50 generations after the last contraction. Finally, we used linear regressions to test the relationship between age and relative fitness, and between age and the number of offspring.

## Results

Simulated range contraction models resulted in population declines of 60–80%, with the largest decreases occurring in *shrinkage* and *amputation* **(Figure 2a).** The model with the lowest population decline (60%) was *hollow*. All models showed significant increases in average relatedness (*p* < 0.001 in all cases; *r*^2^ ranged from 0.33–0.65; **Figure 2b**), but the slope of the relationship between relatedness and generations was steepest in the *shrinkage* and *hollow* models. Variability in relatedness increased following range contraction (**Figure 2b**). While each model showed an eventual decline in nucleotide diversity (π), they differed in the number of generations before π became significantly less than pre-contraction conditions (**Figure 3**). After 25 generations, π was notably reduced in *amputation*, *fragmentation*, and *hollow* simulations, but not in *shrinkage*. In the *shrinkage* models, π did not show marked decline until 100 generations after range contraction. In addition, π was strongly influenced by spatial location, with some models maintaining regions of high absolute and relative π (**Figures S3**, **S7**, **S9**). For all models, age was significantly correlated with number of offspring (*p* < 0.001; *r*^2^ = 0.50; **Figure S1**) and the average and max age of individuals increased as species’ ranges contracted (*p* < 0.001; **Figure 2c–d**). Age was also significantly negatively correlated with fitness, though the fit was poor (*p* < 0.001; *r*^2^ = 0.01).

**Figure 2.**
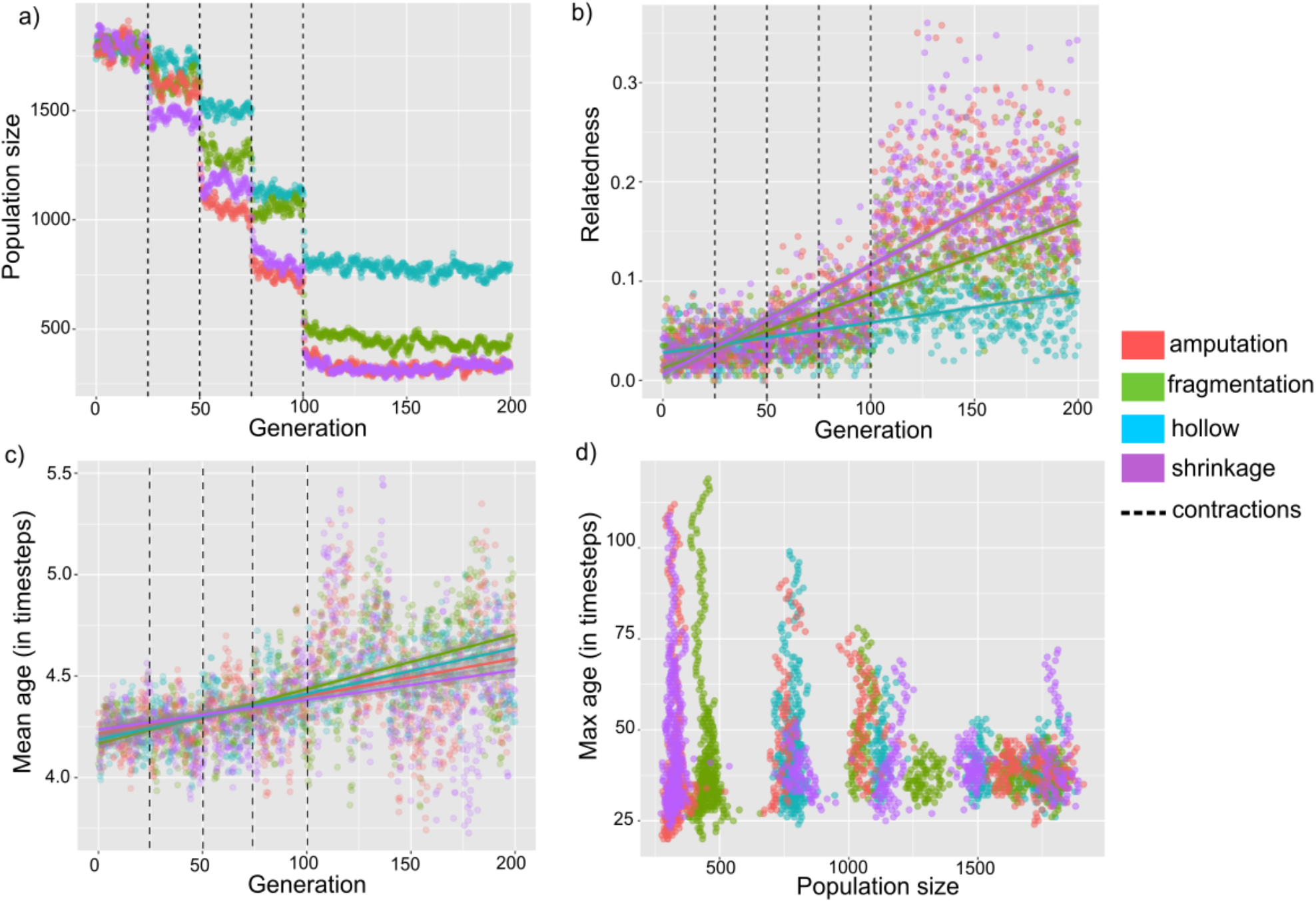
Effects of patterns of range contraction on populations size and demography. a) Population size decreased following each discrete range contraction event; b) increase in pedigree relatedness across generations increased following contraction, with different slopes of the relationship between relatedness and generations for different forms of contraction; c) mean age of individuals in the population increased following range contraction; d) relationship between the maximum age of individuals in the population and the absolute population size revealed decreasing trends in maximum age following range contraction. Dashed lines represent the timing of each discrete range contraction event.

**Figure 3.**
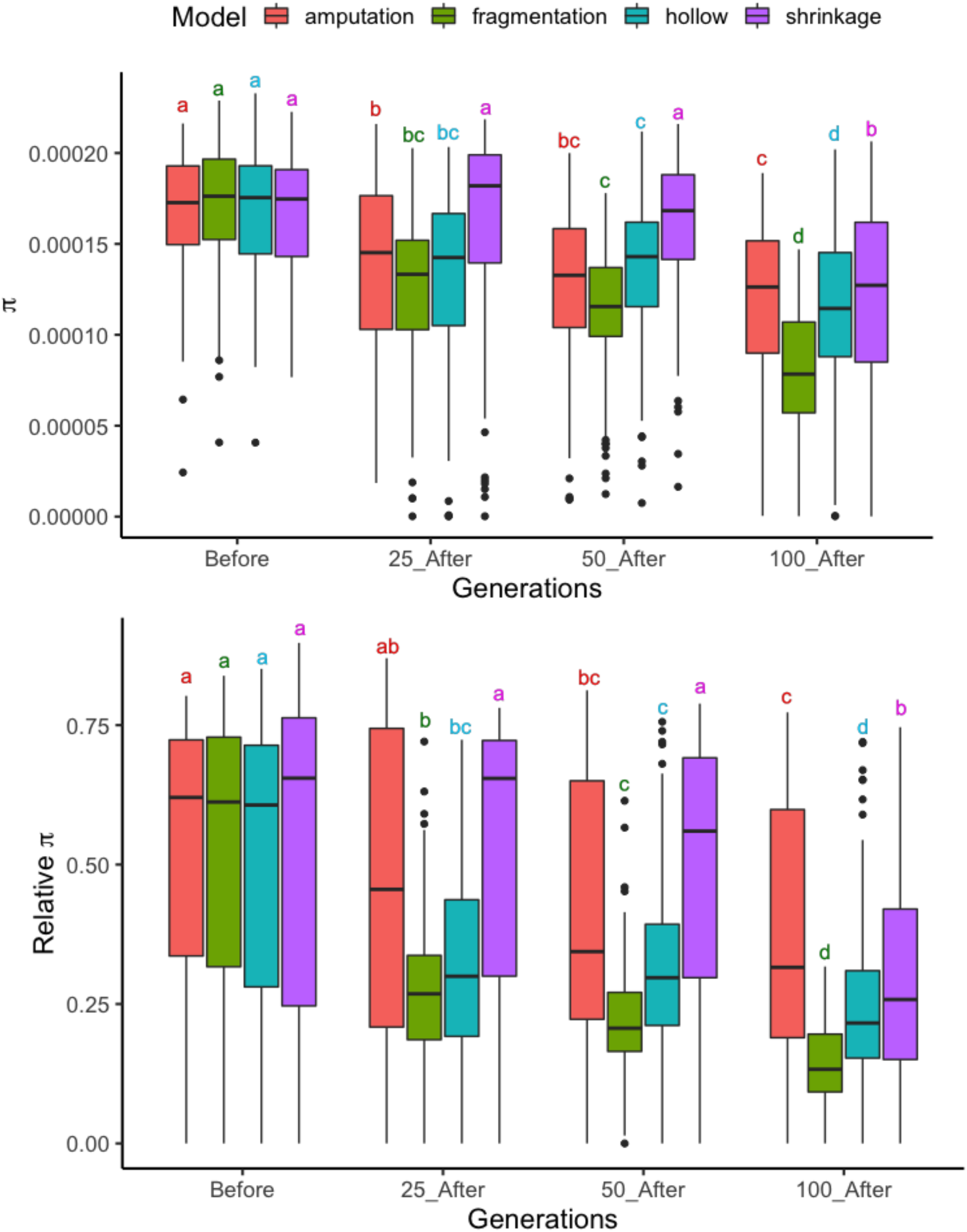
Changes in nucleotide diversity (π) following contraction. a) change in absolute π following the contraction; b) change in π relative to pre-contraction levels. “Before” is 25 generations before the contraction begins; “25_After” is 25 generations after the last contraction point; “50_After” is 50 generations later; and “100_After” is 100 generations later. Tukey’s post hoc test results for differences in π across different time points following each form of range contraction also provided for each plot.

### Shrinkage

The *shrinkage* model resulted in 88% of total range loss, with the remaining area concentrated in the core of the pre-contraction range, as predicted by the demographic hypothesis (Table S1). In the *shrinkage* models, π remained relatively high during and following contraction when compared with the other forms of range contraction. Nucleotide diversity was not significantly different pre- and post-contraction until 100 generations after the contraction ended, despite an 80% decline in population size **(Figure 3)**. Genetic diversity relative to pre-contraction conditions remained comparable for individuals in the center of the post-contraction range even 100 generations later (**Figure S3**). High genetic diversity was coupled with comparably high relative fitness in the center of the range compared to the edges (**Figure S4**). The total branch distance (in number of timesteps) in the center group was 16045 compared to 10970–13531 for the four corners **(Table S3).** While heterozygosity was not significantly different from pre-contraction conditions until well after the contraction ended, the average relatedness was significantly higher within the first 40–60 generations *during* the contraction (*p* < 0.001). Patterns of genetic differentiation indicate spatial autocorrelation of ancestry, with pairwise *F*_ST_ the highest among edge pairs and lowest when edges were compared to the “center” group (**Table S4**). Ancestry was spread evenly across the range, in the *shrinkage* model (**Figure 4**).

**Figure 4.**
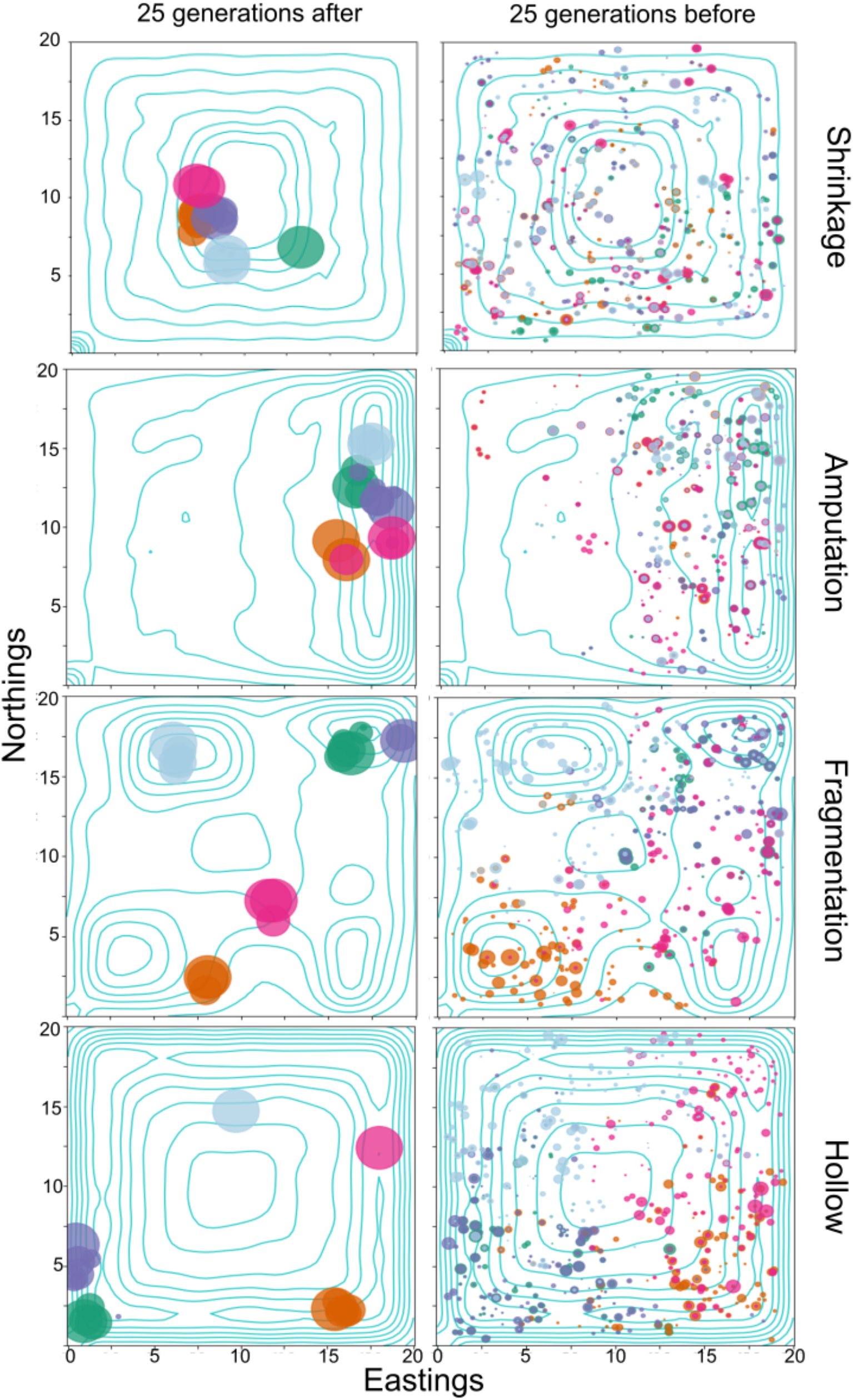
The spread of spatial ancestry of 5 randomly sampled individuals in the post-contraction range. Left column is sampled individuals from the range 25 generations after the contraction. The size of the circles indicate the proportion of the genome inherited from a given ancestor. Note that there are multiple circles due to overlapping generations. Each color represents the sampled individuals. Right column is the ancestors of the 5 sampled individuals 25 generations prior to the contraction.

### Amputation

The *amputation* model resulted in 87% total range loss, with the post-contraction range constrained to the periphery of the pre-contraction range, as consistent with the contagion hypothesis. The highest relative fitness in the *amputation* model was concentrated in the core of the post-contraction range (**Figure S6**), and this area further maintained the highest diversity relative to pre-contraction conditions. By 25 generations post-contraction, π was significantly less than pre-contraction levels (*p* < 0.001). Of the five sampled groups (see **Figure S5, Table S5**), the ‘middle’ group had the highest diversity (π = 1.44e-04), whereas the ‘top’ and ‘lower’ had the lowest (π = 1.11e-04; π = 9.46e-05, respectively). A clear pattern of isolation-by-distance had formed as well, with ‘top’ being least diverged from ‘upper middle’ and most from ‘lower’ (**Table S7**). The spatial ancestry spread was strongly biased in the direction of the extinction force (**Figure 4**).

### Hollow

The *hollow* model resulted in the least amount of range loss of all four patterns, at 68%. The post-contraction range was located in the periphery of the pre-contraction range, consistent with the contagion hypothesis. *Hollow* also resulted in the smallest population size decline of ~60%, which was reflected in a poor fit of average relatedness over generations (*r*^2^ = 0.33). However, *hollow* produced the strongest positive relationship between mean age and generation time (*p* < 0.001; *r*^2^ = 0.43). The *hollow* model demonstrated that pockets of high fitness appeared freely across the range, rather than in the center of the range (**Figure S10**). Sampled groups did not show dramatic differences in π or branch distances (**Table S11–12**). Furthermore, no clear pattern emerged in respect to genetic differentiation with distance (**Table S13**), Spatial ancestry was strongly biased towards the edges, with few ancestors spanning the entire range prior to the contraction (**Figure 4**).

### Fragmentation

The *fragmentation* model resulted in 86% total range loss, with the post-contraction range consisting of disjunct fragments. The *fragmentation* model resulted in the largest impact on genetic diversity. Diversity decreased in a steady stepwise manner after the contraction (**Figure 3**). By 100 generations after the start of the contraction, no sampled individuals maintained π comparable to the pre-contraction mean (**Figure S7**). The four sampled groups were strongly differentiated from one another (*F*_ST_ > 0.42 for all comparisons), and the average branch-distance was twice as deep between groups as within groups (**Table S9–10**). The highest relative fitness was within the center of the remaining groups (see **Figure S8**), decreasing toward the edges. Spatial ancestry was skewed towards the remaining fragment that the sample was taken from, with fragments with the highest population density (‘topleft’ and ‘bottomleft’) showing the greatest spatial spread (**Figure 4**).

## Discussion

Our results demonstrated how range contractions can contribute to biological annihilation through marked impacts on population size, structure, and genetic diversity. Our simulations revealed that the extent and magnitude of the effects of range contraction differed depending on the pattern of contraction. The unique outcomes resulting from each pattern underscore the importance of documenting how range contractions occur in real-world ecosystems. Our models are an important step towards a general understanding of what impacts are likely to manifest under different range contraction patterns. This is crucial when considering conservation strategies for recovery of populations lost to biological annihilation.

### Genetic and demographic consequences of range contraction patterns

The sensitivity of π to reductions in population size has been a topic of debate for some time, (e.g., Ringbauer *et al.,* 2017; Bradburd & Ralph, 2019) particularly whether π responds to population reductions within a few generations, which is often the timescale of relevance to anthropogenic causes of range contractions. Concordant with previous studies (e.g., Barton *et al.,* 2013; Aguillon *et al.,* 2017), we found that average relatedness responded much more rapidly to reductions in absolute population sizes than π for our simulated generalist species. This occurred in all four range contraction patterns. In the most extreme case, *shrinkage*, we found that a decline in π may not be detectable until >100 generations after range contraction despite a >80% loss in suitable habitat (**Figure 3**). This finding demonstrates a pressing need for the field of conservation genetics to adopt more sensitive measures of population health than π, such as length of identity-by-descent (IBD) tracts (see Chen *et al.,* 2019).

A long-standing assumption in conservation genetics is that low genetic diversity (measured as π) increases extinction probability and must result from population bottlenecks (Amos & Balmford, 2001). However, it has been argued that some populations maintain low diversity ancestrally, making this statistic a poor predictor of the population’s health. The cheetah (*Acinonyx jubatus*) and several other species are classic examples of long-term patterns of low genetic diversity without ill effects (O’Brien *et al.,* 1986; Caro & Laurenson, 1994; Caughley, 1994; Merola, 1994; Frankham, 1995; Caro, 2000; Amos & Balmford, 2001).

Our range contraction models show that loss of genetic diversity *relative* to pre-contraction levels may be more indicative of potential impacts on the health of remaining populations. We demonstrated the rate of decline of π within a population was highly impacted by the spatial pattern of contraction. Contraction patterns that maintained high connectivity and impacted the periphery of the population most heavily (such as *shrinkage*) tended to be resilient to declines in π. Because the population density was highest in the core of the range, the loss of peripheral populations did not remove the bulk of standing diversity (Wilkins & Wakeley, 2002). As expected, the contractions that constrained the remaining range towards the edges, as in *hollow* and *amputation*, caused appreciable reductions in standing diversity despite maintaining absolute population sizes similar to *shrinkage*. Furthermore, the loss of connectivity (as in *fragmentation*) had dramatic impacts on the rate of decline of π. Reduced connectivity has been recognized as an important driver in extinction risk of populations (Keller *et al.,* 2003; Chan *et al.,* 2020).

Discrete sampling in continuous populations is known to bias measures of dispersal and connectivity (Wang & Bradburd, 2014; Cayuela *et al.,* 2018; Battey *et al.,* 2020). This is partially due to the metrics of gene flow (such as *F*_ST_) being derived for discrete populations. Furthermore, incomplete sampling across the range may skew the interpretation of the impact of a contraction on measures of diversity. In our simulations, we found that samples taken from the range center consistently had higher π and lower differentiation than those from the edges (**Tables S2–3, S5–6**). Indeed, for the *shrinkage* pattern, the level of π in the range center was comparable to the pre-contraction conditions long after the contraction ended. This demonstrates the importance of having prior knowledge about range size and boundaries for species of interest and species distributions throughout the range.

Range contractions also contribute to biological annihilation by altering demographics of populations. Indeed, some alterations in demographic patterns are expected to become apparent following shifts in absolute population size. For example, the age structure of a population has been found to shift towards older age classes following population declines, which has been attributed to reduced survival of juveniles or reproductive failure (Stubbs & Swingland, 1985; Wheeler *et al.,* 2003). Our models produced the same trends (**Figure 2c–d**). However, our models have no age-specific fitness declines; in effect, these correlations emerged as a by-product of the increased stochasticity of smaller populations. We recommend that future investigations into the age structure of declining populations account for this increased randomness when providing explanations for shifts towards older age classes. This will likely include an experimental or observational approach that extends beyond counts of individuals per age class to more mechanistic, instead of inferential, causes of shifts.

### Implications for Theories of Geographic Range

Channell and Lomolino (2000*a, b*) found that, with a few exceptions, range contractions consistent with the contagion hypothesis (i.e., *amputation* and *hollow*) were far more likely to occur than those predicted by the demographic hypothesis. As such, our generalizations for *amputation* and *hollow* contagion patterns will likely be the most broadly applicable in natural systems, though the *hollow* pattern was mostly theoretical in this analysis.

Though *amputation* may be the more common contagion pattern of range contraction in natural systems (as opposed to *hollow*), our findings reveal that peripheral populations will be differentially impacted depending on the way the contagion spreads. Whereas the range center maintained the highest relative fitness in the *amputation* pattern, the *hollow* range demonstrated that pockets of high fitness appeared to shift freely across the range (**Figure S10**). This may be a response to shifting population density: when the density becomes too high in any one area, relative fitness declines causing the highest fitness to shift away from the area. However, these areas are somewhat skewed towards corners, which may be due to the ability of corners to maintain higher population densities than straights due to the greater proximity of habitable areas (well within the dispersal distance). Pairwise *F*_ST_ among sampled groups was relatively high (0.27–0.37) in *hollow* compared to *amputation*. This difference, and the greater decline in relative π, may be the result of higher population densities in the *amputation* model relative to the *hollow* model. Spatial ancestry in the *hollow* simulation was strongly biased towards the edges, with few ancestors spanning the entire range prior to the contraction, as opposed to the *amputation* simulation, where spatial ancestry was biased in the direction of the extinction force (**Figure 4**). Given that the importance of the range periphery relative to range core for species persistence has been contested in the literature (Channell, 2004; Channell & Lomolino, 2000*a*,*b*; Rodríguez, 2002), we caution that peripherally contracted ranges will likely vary according to how extinction factors spread across each range.

*Fragmentation* is a ubiquitous and challenging form of range contraction and biological annihilation (Fitzgerald *et al.*, 2018), yet it has not been adequately addressed in studies of geographic range (but see Donald & Greenwood, 2001). Our simulations of range contraction by *fragmentation* resulted in more drastic effects on genetic diversity and post-contraction population genetic structure than the other patterns. Range fragmentation can occur naturally over geologic time scales, yet is also caused by human land use over rapid time scales (Chan *et al.,* 2020). Range fragmentation has also been shown to cause striking demographic disruption (Cegelski *et al.,* 2003; Proctor *et al.,* 2005) that in some instances has directly led to population extinction (Leavitt & Fitzgerald, 2013; Walkup *et al.,* 2017). Most fragmentation research is directed at understanding effects of habitat fragmentation on populations (Fahrig, 2003; Rogan & Lacher, 2018). Including *fragmentation* in geographic range theory is especially relevant considering that land use occurs across multiple temporal and spatial scales and has driven spatial patterns of range contraction (Boakes *et al.,* 2018). We suggest that *fragmentation* merits further consideration as an important pattern of range contraction across the globe.

### Future Research and Implications for Conservation

The principal implication of our results is that a “one size fits all” conservation approach will not be effective in ameliorating the consequences of range contraction. We showed *fragmentation* caused strong genetic differentiation among disjunct range fragments (*F*_ST_ > 0.42 for all comparisons), which resulted in increased pedigree relatedness and decreased genetic diversity in each range fragment due to inbreeding. However, genetic diversity as whole remained high because the action of genetic drift was partitioned to individual fragments. This finding supports conservation corridor theory and conservation translocations in the form of population reinforcement. In natural systems, it may be a priority to develop corridors between fragments to restore gene flow and employ reciprocal introductions to mitigate loss of diversity among remnant populations (Hurtado *et al.* 2012). Reintroductions may be an important strategy for the *amputation* scenario, in which connectivity remained high in the remaining range but genetic diversity was low due to the persistence of historically less diverse lineages. Undoubtedly, a complex synergy of unique factors including life history, phylogeny, social group structure, behavioral flexibility, ecological niche, or local and regional factors (Lawton, 1993; Purvis *et al.,* 2000; Cardillo *et al.,* 2008; Boakes *et al*., 2018) should be considered when developing strategies to combat biological annihilation.

Our finding that the spatial distribution of ancestors was strongly skewed in the direction of the contraction force for almost all patterns bears important implications for interventions aimed at addressing the loss of locally adapted gene complexes. This ancestry effect was particularly pronounced in *amputation,* the most common form of contraction. Spatial ancestry was influenced in our models by local lineages more than through migration. This implies that local adaptations may be lost because lineages carrying those adaptations go extinct as the range contracts. Attempts at repatriation in the historic range may be hindered by lack of locally adapted gene complexes, and conservation interventions should be designed to monitor and prevent loss of local lineages (Templeton, 1986).

The timing and rate of response of π and demographic factors varied among the four patterns we investigated. We found that pedigree relatedness in *shrinkage* models increased well ahead of the decline in π. Time lags between range contraction and detectable differences in genetic diversity has been observed in several other studies (Anderson *et al.,* 2010; Balkenhol *et al*., 2016; Schlaepfer *et al.,* 2018). We therefore suggest measures other than π such as relatedness may generally show a quicker response and should be considered when assessing impacts of range contractions. Furthermore, the importance of recognizing early changes in demographic structure based on the pattern of range contraction may increase capacity to rescue populations before they are severely compromised.

Our simulation model has a few limitations to consider. First, individuals in our models are hermaphroditic, which alleviates the issue in small populations of finding a mate of the opposite sex. Thus, our results represent a conservative measure of the impacts of range contraction. Future work might consider modelling separate sexes and more complex mating systems. Second, despite living for several generations, individuals only disperse once immediately after birth, which limits their ability to respond to range contractions. For highly vagile organisms that may reproduce in several locations over their lifetimes, our results will be exaggerated. This limitation can also be mitigated by future work incorporating adult movement following offspring generation, which will allow a greater number of individuals to “escape” the extinction front. Thirdly, our simulated ranges are uniform in their pre-contraction suitability, whereas natural ranges are typically patchier. Finally, we assume that contraction happens in discrete intervals instead of continuously. We do this for model simplicity, but we recognize that some contractions may happen continuously, depending on the driver(s).

Empirical studies that explicitly address range contraction patterns are of increasing value to conservation, especially if genetic and demographic correlates are also measured. Patterns of range contraction have typically only been considered in multi-species analyses and reviews (e.g., Laliberte & Ripple, 2004), while most reports of range contractions for single species focus on the amount and extent of range lost. (however, see Lomolino & Smith, 2001). Our results show an important next step will be to investigate consequences of contraction patterns, particularly *fragmentation*, in real ecological systems. Understanding range contraction patterns and their consequences for the planet’s biodiversity is crucial to further combat biological annihilation in the Anthropocene.

## Acknowledgements

Special thanks to Peter Ralph and Ben Haller for coding help. We thank the Applied Biodiversity Science Program weekly journal club where the idea for this paper originated.

## SUPPORTING INFORMATION

Additional Supporting Information may be downloaded via the online version of this article at Wiley Online Library (www.ecologyletters.com).

#### Box 1. Range contractions: hypotheses and patterns

The mechanisms of range contraction have been predominately described by two hypotheses: *demographic* and *contagion* (Lomolino and Channell,1995; Channell & Lomolino, 2000*b*). The *demographic hypothesis* states that local populations go extinct because of demographic processes in increasingly small populations. It is derived from the theory of extinction in small populations and is predicated on the assumption that populations are larger and denser in the core of a species range. Therefore, the demographic hypothesis predicts that all else being equal, populations should go extinct beginning on the periphery of the range, and the last remaining populations will be found closest to the center of the species’ historical range (Channell and Lomolino 2000*b*). The demographic hypothesis does not consider drivers of extinction.

The *contagion hypothesis,* which was favored by Lomolino and Channell (1995, 1998), posits that multiple extinction factors (e.g., habitat loss, climate change, disease, persecution) take place in concert across landscapes and these extinction factors spread like a contagion causing areas of the range to be lost. The contagion hypothesis predicts that if complete extinction does not occur, remaining populations will persist in the periphery of the range where they are most geographically isolated from the extinction factors (Towns and Daugherty 1994, Lomolino and Channel 1995). By examining the changes in the index of centrality of contracting ranges, Channell and Lomolino (2000*b*) found that initial range loss was usually peripheral. However, the initial impact of extinction factors does not have to occur in the periphery to be consistent with the contagion hypothesis (Channell and Lomolino 2000b, Donald and Greenwood 2001).

Here we considered four patterns of species range contraction and the potential mechanisms that form them: *shrinkage, amputation*, *hollow*, and *fragmentation*. *Shrinkage* is the predicted outcome from the demographic hypothesis and describes the scenario where a species’ range contracts from its periphery to its core. This pattern has also been referred to as a “melting range” (Rodríguez, 2002) or “range collapse” (Fisher, 2011). *Amputation* is predicted by the contagion hypothesis and occurs when loss of geographic range begins at a particular point or region in the periphery of the species’ range and spreads across the range until the last remaining populations occur in the areas that are furthest from the original population extinctions. In short, portions of the species’ range are amputated with the advancing extinction front. This is thought to be the most common way that ranges contract (Lomolino and Channel 1995, Channel and Lomolino 2000*a*,*b*). *Hollow* is also derived from the contagion hypothesis, yet distinct from range amputation in that extinction begins in the center of a species’ range and spreads until the remaining populations only persist on the periphery. *Fragmentation* occurs through loss of continuity in a species’ range, resulting in disjunct populations, and constrains dispersal, contributing to impacts on genetic diversity. *Fragmentation* may result in a much smaller total area occupied by a species even if the geographic span of the range is similar to its historical baseline. Range contraction through *fragmentation* does not conform to either the demographic or contagion hypothesis. Though pattern is often not considered in case studies of range contractions, it can be inferred, especially when contractions are recent and historical range data exists. With the exception of *hollow*, which is mostly theoretical in our study, range contractions demonstrating these patterns have been observed in a wide variety of taxa worldwide and have been attributed to a number of different drivers (**Table 1**).

## Notes

### Competing Interest Statement

The authors have declared no competing interest.

