## Supplementary Material for "Paths to annihilation: Genetic and demographic consequences of range contraction patterns"

**Tables**

Table S1. Summary of impacts of forms of range contraction in the post-contraction period relative to the pre-contraction population.

| **Model** | **Percent range lost** | **Relative fitness** | **Population density** | **Genetic diversity (π)** | **Spatial ancestry** |
| --- | --- | --- | --- | --- | --- |
| *Shrinkage* | 88% | Highest in the center, decreases towards edges | Concentrated in the center of the range | Maintains high relative and absolute π | Approximates random distribution |
| *Amputation* | 87% | Highest in center, decreases north-south | Spread thinly across the range | Lower overall; relative π around mean in the center | Strongly skewed in the direction of the extinction front |
| *Hollow* | 68% | Shifts around the range, consistently highest near corners | Highest in the corners, decreasing along straights | Lower overall; some pockets of relative π near mean | Skewed toward range edges |
| *Fragmentation* | 86% | Highest around surviving deme centers, decreases along edges | Concentrated in center of surviving demes | Lowest overall; no individuals maintain π near mean | Skewed towards location of surviving deme; higher density demes with greater spatial spread |

Table S2. Pairwise comparisons of π_12_ values for sampled groups in the *shrinkage* simulation models. In each quadrant and center of the range, and in the pre-contraction sample (“ancient”), 50 individuals were sampled.

|  | Topleft | Topright | Bottomleft | Bottomright | Center | Ancient |
| --- | --- | --- | --- | --- | --- | --- |
| Topleft | 0.0001098 |  |  |  |  |  |
| Topright | 0.0001816 | 0.0001219 |  |  |  |  |
| Bottomleft | 0.0001886 | 0.000186 | 0.0001351 |  |  |  |
| Bottomright | 0.0001886 | 0.0001696 | 0.0001639 | 0.000129 |  |  |
| Center | 0.0001774 | 0.0001678 | 0.0001676 | 0.0001604 | 0.0001603 |  |
| Ancient | 0.0002145 | 0.0002195 | 0.0002161 | 0.0002152 | 0.0002168 | 0.0002057 |

Table S3. Pairwise comparisons of branch distance values for sampled groups in the *shrinkage* simulation models. In each quadrant and center of the range, and in the pre-contraction sample (“ancient”), 50 individuals were sampled.

|  | Topleft | Topright | Bottomleft | Bottomright | Center | Ancient |
| --- | --- | --- | --- | --- | --- | --- |
| Topleft | 10970.1326 |  |  |  |  |  |
| Topright | 18149.0073 | 12194.3461 |  |  |  |  |
| Bottomleft | 18859.433 | 18616.6526 | 13531.94 |  |  |  |
| Bottomright | 18866.9304 | 16970.4375 | 16406.1384 | 12907.6828 |  |  |
| Center | 17736.3461 | 16791.5995 | 16781.3777 | 16059.387 | 16045.8891 |  |
| Ancient | 21457.2603 | 21969.62 | 21651.3032 | 21543.6693 | 21711.3687 | 20590.4148 |

Table S4. Pairwise comparisons of *F_ST_* values for sampled groups in the *shrinkage* simulation models. In each quadrant and center of the range, and in the pre-contraction sample (“ancient”), 50 individuals were sampled.

|  | Topleft | Topright | Bottomleft | Bottomright | Center | Ancient |
| --- | --- | --- | --- | --- | --- | --- |
| Topleft |  |  |  |  |  |  |
| Topright | 0.221 |  |  |  |  |  |
| Bottomleft | 0.213 | 0.183 |  |  |  |  |
| Bottomright | 0.225 | 0.150 | 0.107 |  |  |  |
| Center | 0.135 | 0.087 | 0.063 | 0.052 |  |  |
| Ancient | 0.152 | 0.145 | 0.118 | 0.125 | 0.084 |  |

Table S5. Pairwise comparisons of π_12_ values for sampled groups in the *amputation* simulation models. In each quadrant and center of the range, and in the pre-contraction sample (“ancient”), 50 individuals were sampled.

|  | Top | Uppermiddle | Middle | Lowermiddle | Lower | Ancient |
| --- | --- | --- | --- | --- | --- | --- |
| Top | 1.11e-04 |  |  |  |  |  |
| Uppermiddle | 1.50e-04 | 1.40e-04 |  |  |  |  |
| Middle | 1.88e-04 | 1.70e-04 | 1.44e-04 |  |  |  |
| Lowermiddle | 2.07e-04 | 1.99e-04 | 1.70e-04 | 1.37e-04 |  |  |
| Lower | 2.07e-04 | 2.01e-04 | 1.76e-04 | 1.44e-04 | 9.46e-05 |  |
| Ancient | 2.18e-04 | 2.18e-04 | 2.17e-04 | 2.16e-04 | 2.16e-04 | 2.08e-04 |

Table S6. Pairwise comparisons of branch distance values for sampled groups in the *amputation* simulation models. In each quadrant and center of the range, and in the pre-contraction sample (“ancient”), 50 individuals were sampled.

|  | Top | Uppermiddle | Middle | Lowermiddle | Lower | Ancient |
| --- | --- | --- | --- | --- | --- | --- |
| Top | 11110 |  |  |  |  |  |
| Uppermiddle | 14975 | 14004 |  |  |  |  |
| Middle | 18778 | 16986 | 14356 |  |  |  |
| Lowermiddle | 20714 | 19869 | 17019 | 13740 |  |  |
| Lower | 20682 | 20049 | 17659 | 14412 | 9494 |  |
| Ancient | 21731 | 21771 | 21729 | 21645 | 21594 | 20778 |

Table S7. Pairwise comparisons of *F_ST_* values for sampled groups in the *amputation* simulation models. In each quadrant and center of the range, and in the pre-contraction sample (“ancient”), 50 individuals were sampled.

|  | Top | Uppermiddle | Middle | Lowermiddle | Lower | Ancient |
| --- | --- | --- | --- | --- | --- | --- |
| Top |  |  |  |  |  |  |
| Uppermiddle | 0.0882 |  |  |  |  |  |
| Middle | 0.192 | 0.090 |  |  |  |  |
| Lowermiddle | 0.251 | 0.178 | 0.096 |  |  |  |
| Lower | 0.336 | 0.261 | 0.194 | 0.107 |  |  |
| Ancient | 0.154 | 0.112 | 0.106 | 0.112 | 0.176 |  |

Table S8. Pairwise comparisons of π_12_ values for sampled groups in the *fragmentation* simulation models. In each quadrant and center of the range, and in the pre-contraction sample (“ancient”), 50 individuals were sampled.

|  | Topleft | Topright | Bottomleft | Bottomright | Ancient |
| --- | --- | --- | --- | --- | --- |
| Topleft | 9.11e-05 |  |  |  |  |
| Topright | 2.21e-04 | 8.55e-05 |  |  |  |
| Bottomleft | 2.29e-04 | 2.34e-04 | 8.26e-05 |  |  |
| Bottomright | 2.33e-04 | 2.18e-04 | 2.27e-04 | 7.08e-05 |  |
| Ancient | 2.15-04 | 2.17e-04 | 2.17e-04 | 2.19e-04 | 2.09e-04 |

Table S9. Pairwise comparisons of branch distance values for sampled groups in the *fragmentation* simulation models. In each quadrant and center of the range, and in the pre-contraction sample (“ancient”), 50 individuals were sampled.

|  | Topleft | Topright | Bottomleft | Bottomright | Ancient |
| --- | --- | --- | --- | --- | --- |
| Topleft | 9124 |  |  |  |  |
| Topright | 22139 | 8595 |  |  |  |
| Bottomleft | 22887 | 23457 | 8273 |  |  |
| Bottomright | 23361 | 21898 | 22697 | 7068 |  |
| Ancient | 21514 | 21748 | 21737 | 21859 | 20950 |

Table S10. Pairwise comparisons of *F_ST_* values for sampled groups in the *fragmentation* simulation models. In each quadrant and center of the range, and in the pre-contraction sample (“ancient”), 50 individuals were sampled.

|  | Topleft | Topright | Bottomleft | Bottomright | Ancient |
| --- | --- | --- | --- | --- | --- |
| Topleft |  |  |  |  |  |
| Topright | 0.429 |  |  |  |  |
| Bottomleft | 0.449 | 0.471 |  |  |  |
| Bottomright | 0.484 | 0.473 | 0.494 |  |  |
| Ancient | 0.177 | 0.191 | 0.196 | 0.219 |  |

Table S11. Pairwise comparisons of π_12_ values for sampled groups in the *hollow* simulation models. In each quadrant and center of the range, and in the pre-contraction sample (“ancient”), 50 individuals were sampled.

|  | Topleft | Topright | Bottomleft | Bottomright | Ancient |
| --- | --- | --- | --- | --- | --- |
| Topleft | 1.18e-04 |  |  |  |  |
| Topright | 2.14e-04 | 9.71e-05 |  |  |  |
| Bottomleft | 2.19e-04 | 2.31e-04 | 1.29e-04 |  |  |
| Bottomright | 2.28e-04 | 2.13e-04 | 2.18e-04 | 9.48e-05 |  |
| Ancient | 2.13e-04 | 2.15e-04 | 2.18e-04 | 2.16e-04 | 2.08e-04 |

Table S12. Pairwise comparisons of π_12_ values for sampled groups in the *hollow* simulation models. In each quadrant and center of the range, and in the pre-contraction sample (“ancient”), 50 individuals were sampled.

|  | Topleft | Topright | Bottomleft | Bottomright | Ancient |
| --- | --- | --- | --- | --- | --- |
| Topleft | 11800 |  |  |  |  |
| Topright | 21397 | 9695 |  |  |  |
| Bottomleft | 21848 | 23102 | 12864 |  |  |
| Bottomright | 22820 | 21356 | 21785 | 9504 |  |
| Ancient | 21273 | 21529 | 21811 | 21558 | 20811 |

Table S13. Pairwise comparisons of *F_ST_* values for sampled groups in the *fragmentation* simulation models. In each quadrant and center of the range, and in the pre-contraction sample (“ancient”), 50 individuals were sampled.

|  | Topleft | Topright | Bottomleft | Bottomright | Ancient |
| --- | --- | --- | --- | --- | --- |
| Topleft |  |  |  |  |  |
| Topright | 0.331 |  |  |  |  |
| Bottomleft | 0.278 | 0.343 |  |  |  |
| Bottomright | 0.364 | 0.379 | 0.321 |  |  |
| Ancient | 0.132 | 0.171 | 0.129 | 0.174 |  |

Table S14. Results of the Tukey’s post hoc test for differences in π across different time points following range contraction for the *shrinkage* model.

| Comparison | Difference | *p*-value |
| --- | --- | --- |
| Before – 25 generations after | 6.52e-05 | 0.774 |
| Before – 50 generations after | 8.70e-06 | 0.579 |
| Before – 100 generations after | 4.94e-05 | **0.000** |
| 25 – 50 generations after | 2.17e-06 | 0.988 |
| 25 – 100 generations after | 4.28e-05 | **0.000** |
| 50 – 100 generations after | 4.07e-05 | **0.000** |

Table S15. Results of the Tukey’s post hoc test for differences in relative π across different time points following range contraction for the *shrinkage* model.

| Comparison | Difference | *p*-value |
| --- | --- | --- |
| Before – 25 generations after | 0.016 | 0.962 |
| Before – 50 generations after | 0.048 | 0.493 |
| Before – 100 generations after | 0.238 | **0.000** |
| 25 – 50 generations after | 0.032 | 0.791 |
| 25 – 100 generations after | 0.221 | **0.000** |
| 50 – 100 generations after | 0.189 | **0.001** |

Table S16. Results of the Tukey’s post hoc test for differences in π across different time points following range contraction for the *amputation* model.

| Comparison | Difference | *p*-value |
| --- | --- | --- |
| Before – 25 generations after | 2.69e-05 | **0.001** |
| Before – 50 generations after | 3.71e-05 | **0.000** |
| Before – 100 generations after | 4.65e-05 | **0.000** |
| 25 – 50 generations after | 1.01e-05 | 0.335 |
| 25 – 100 generations after | 1.95e-05 | **0.007** |
| 50 – 100 generations after | 9.44e-06 | 0.394 |

Table S17. Results of the Tukey’s post hoc test for differences in relative π across different time points following range contraction for the *amputation* model.

| Comparison | Difference | *p*-value |
| --- | --- | --- |
| Before – 25 generations after | 0.045 | 0.555 |
| Before – 50 generations after | 0.103 | **0.014** |
| Before – 100 generations after | 0.149 | **0.001** |
| 25 – 50 generations after | 0.058 | 0.319 |
| 25 – 100 generations after | 0.105 | **0.011** |
| 50 – 100 generations after | 0.047 | 0.508 |

Table S18. Results of the Tukey’s post hoc test for differences in π across different time points following range contraction for the *fragmentation* model.

| Comparison | Difference | *p*-value |
| --- | --- | --- |
| Before – 25 generations after | 4.63e-05 | **0.000** |
| Before – 50 generations after | 5.75e-05 | **0.000** |
| Before – 100 generations after | 9.14e-05 | **0.000** |
| 25 – 50 generations after | 1.12e-05 | 0.148 |
| 25 – 100 generations after | 4.50e-05 | **0.000** |
| 50 – 100 generations afte | 3.39e-05 | **0.000** |

Table S19. Results of the Tukey’s post hoc test for differences in relative π across different time points following range contraction for the *fragmentation* model.

| Comparison | Difference | *p*-value |
| --- | --- | --- |
| Before – 25 generations after | 0.254 | **0.000** |
| Before – 50 generations after | 0.311 | **0.000** |
| Before – 100 generations after | 0.388 | **0.000** |
| 25 – 50 generations after | 0.057 | **0.041** |
| 25 – 100 generations after | 0.133 | **0.000** |
| 50 – 100 generations after | 0.078 | **0.001** |

Table S20. Results of the Tukey’s post hoc test for differences in π across different time points following range contraction for the *hollow* model.

| Comparison | Difference | *p*-value |
| --- | --- | --- |
| Before – 25 generations after | 3.88e-05 | **0.000** |
| Before – 50 generations after | 3.37e-05 | **0.001** |
| Before – 100 generations after | 5.52e-05 | **0.000** |
| 25 – 50 generations after | -5.07e-06 | 0.822 |
| 25 – 100 generations after | 1.64e-05 | **0.027** |
| 50 – 100 generations after | 2.15e-05 | **0.002** |

Table S21. Results of the Tukey’s post hoc test for differences in relative π across different time points following range contraction for the *hollow* model.

| Comparison | Difference | *p*-value |
| --- | --- | --- |
| Before – 25 generations after | 0.199 | **0.000** |
| Before – 50 generations after | 0.194 | **0.000** |
| Before – 100 generations after | 0.273 | **0.000** |
| 25 – 50 generations after | -0.005 | 0.998 |
| 25 – 100 generations after | 0.074 | **0.031** |
| 50 – 100 generations after | 0.079 | **0.018** |

Table S22. Mean, standard deviation, and range for π in all contraction models and time points analyzed in Tables S14–21.

| Model | Time point | Mean | St. Dev. | Range |
| --- | --- | --- | --- | --- |
| Shrinkage | Before | 1.17e-04 | 3.45e-05 | 7.67e-05, 2.22e-4 |
| Shrinkage | 25 generations after | 1.16e-04 | 5.54e-05 | 1.79e-07, 2.19e-04 |
| Shrinkage | 50 generations after | 1.16e-04 | 4.32e-05 | 1.64e-05, 2.16e-04 |
| Shrinkage | 100 generations after | 1.12e-04 | 5.65e-05 | 1.09e-07, 2.06e-04 |
| Amputation | Before | 1.65e-04 | 3.60e-05 | 2.43e-05, 2.16e-04 |
| Amputation | 25 generations after | 1.39e-04 | 4.86e-05 | 1.85e-05, 2.16e-04 |
| Amputation | 50 generations after | 1.28e-04 | 4.11e-05 | 9.30e-06, 2.00e-04 |
| Amputation | 100 generations after | 1.18e-04 | 4.29e-05 | 4.04e-07, 1.89e-04 |
| Fragmentation | Before | 1.69e-04 | 3.73e-05 | 4.08e-05, 2.28e-04 |
| Fragmentation | 25 generations after | 1.23e-04 | 4.11e-05 | 1.69e-07, 2.03e-04 |
| Fragmentation | 50 generations after | 1.12e-04 | 3.59e-05 | 1.24e-05, 1.78e-04 |
| Fragmentation | 100 generations after | 7.83e-05 | 3.44e-05 | 2.39e-07, 1.47e-04 |
| Hollow | Before | 1.69e-04 | 3.51e-05 | 4.07e-05, 2.32e-04 |
| Hollow | 25 generations after | 1.31e-04 | 4.61e-05 | 1.43e-07, 2.03e-04 |
| Hollow | 50 generations after | 1.36e-04 | 4.09e-05 | 7.47e-06, 2.12e-04 |
| Hollow | 100 generations after | 1.14e-04 | 4.24e-05 | 2.76e-07, 2.02e-04 |

Table S23. Mean, standard deviation, and range for relative π in all contraction models and time points analyzed in Tables S14–21.

| Model | Time point | Mean | St. Dev. | Range |
| --- | --- | --- | --- | --- |
| Shrinkage | Before | 0.533 | 0.278 | 0.000, 0.898 |
| Shrinkage | 25 generations after | 0.517 | 0.250 | 0.000, 0.781 |
| Shrinkage | 50 generations after | 0.485 | 0.233 | 0.000, 0.789 |
| Shrinkage | 100 generations after | 0.295 | 0.202 | 0.000, 0.746 |
| Amputation | Before | 0.518 | 0.224 | 0.000, 0.803 |
| Amputation | 25 generations after | 0.474 | 0.282 | 0.000, 0.870 |
| Amputation | 50 generations after | 0.416 | 0.231 | 0.000, 0.813 |
| Amputation | 100 generations after | 0.369 | 0.216 | 0.000, 0.773 |
| Fragmentation | Before | 0.527 | 0.235 | 0.000, 0.839 |
| Fragmentation | 25 generations after | 0.273 | 0.137 | 0.000, 0.721 |
| Fragmentation | 50 generations after | 0.217 | 0.108 | 0.000, 0.614 |
| Fragmentation | 100 generations after | 0.139 | 0.070 | 0.000, 0.317 |
| Hollow | Before | 0.524 | 0.231 | 0.000, 0.851 |
| Hollow | 25 generations after | 0.325 | 0.186 | 0.000, 0.724 |
| Hollow | 50 generations after | 0.330 | 0.180 | 0.000, 0.756 |
| Hollow | 100 generations after | 0.251 | 0.155 | 0.000, 0.720 |

**Figures**

Figure S1. Relationship between mean number offspring and mean age for all simulated models.

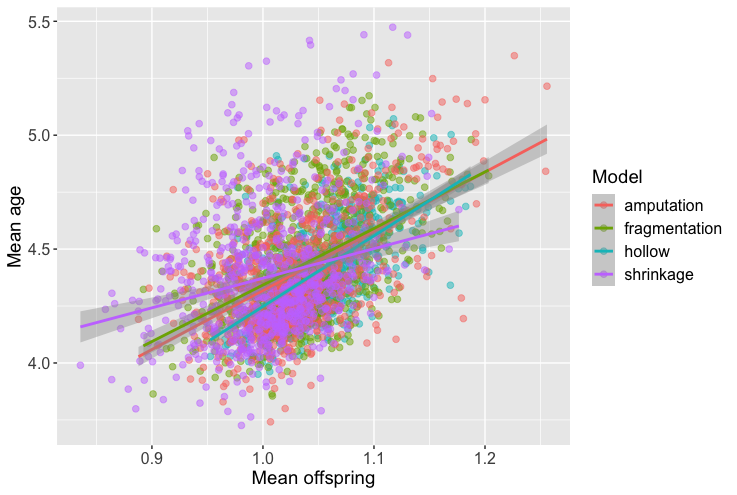

Figure S2. Relationship between average pedigree relatedness and log(population size) for all simulated models.

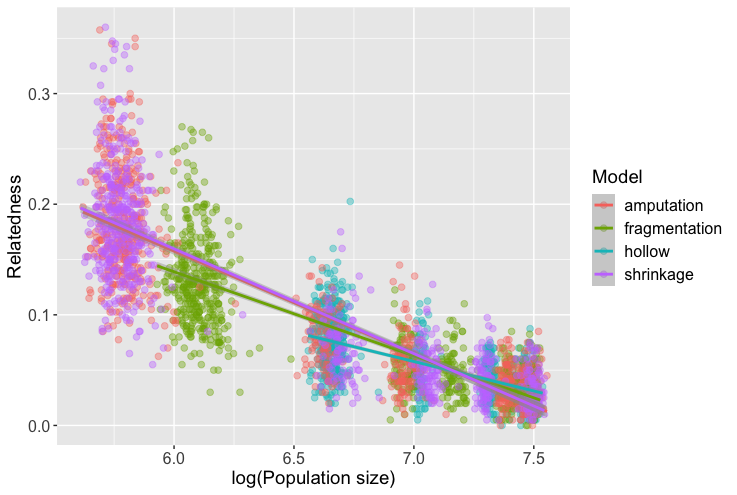

Figure S3. Spatial patterns of π in the *shrinkage* model before the contraction and 25, 50, and 100 generations after. Relative π is scaled to the pre-contraction levels, with values ~0.5 representing the mean. Size of the circle is scaled to absolute π.

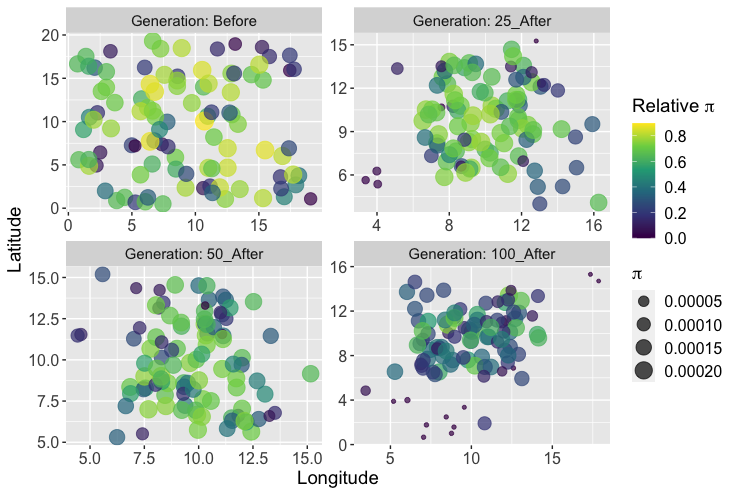

Figure S4. Shifts in relative fitness in the *shrinkage* models at generations 200, 150, 100, and 0.

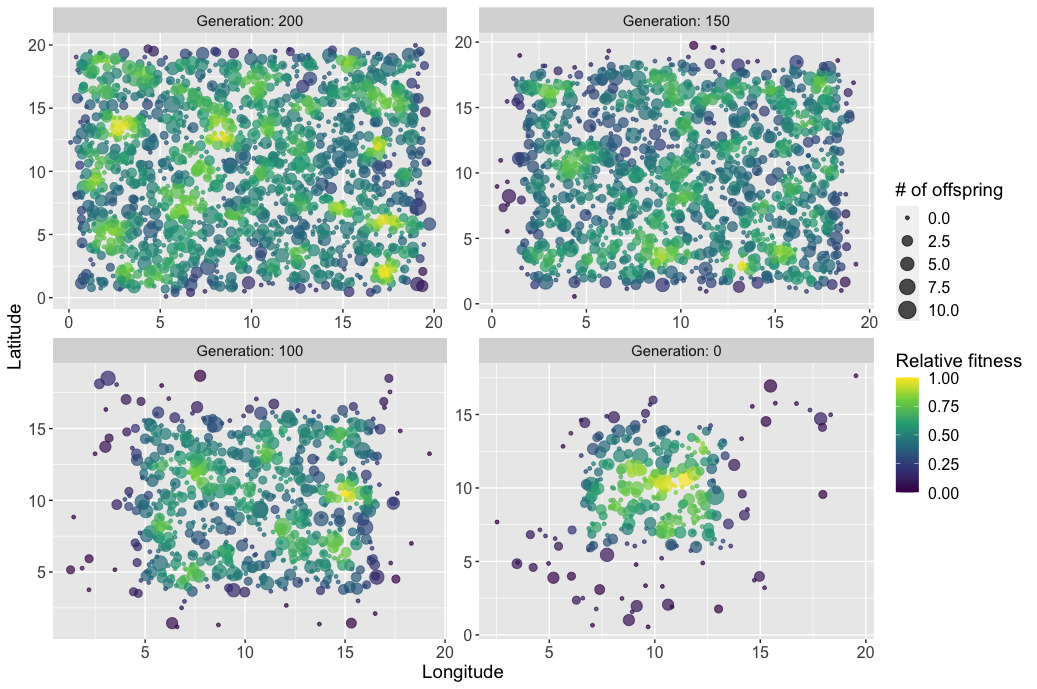

Figure S5. Spatial patterns of π in the *amputation* model before the contraction and 25, 50, and 100 generations after. Relative π is scaled to the pre-contraction levels, with values ~0.5 representing the mean. Size of the circle is scaled to absolute π.

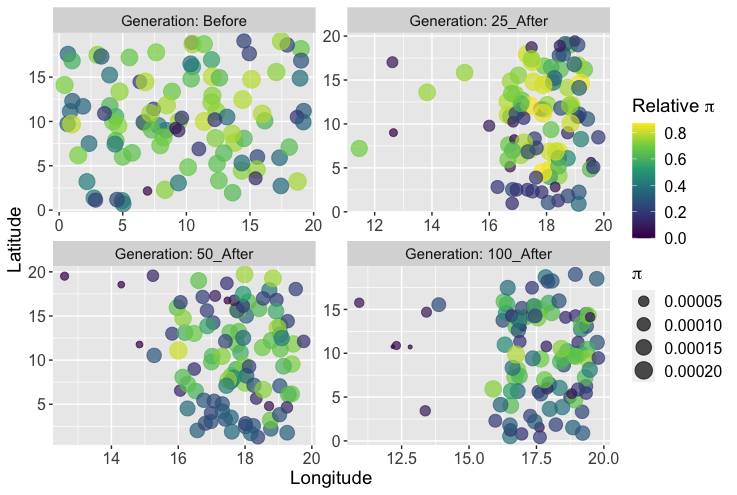

Figure S6. Shifts in relative fitness in the *amputation* models at generations 200, 150, 100, and 0.

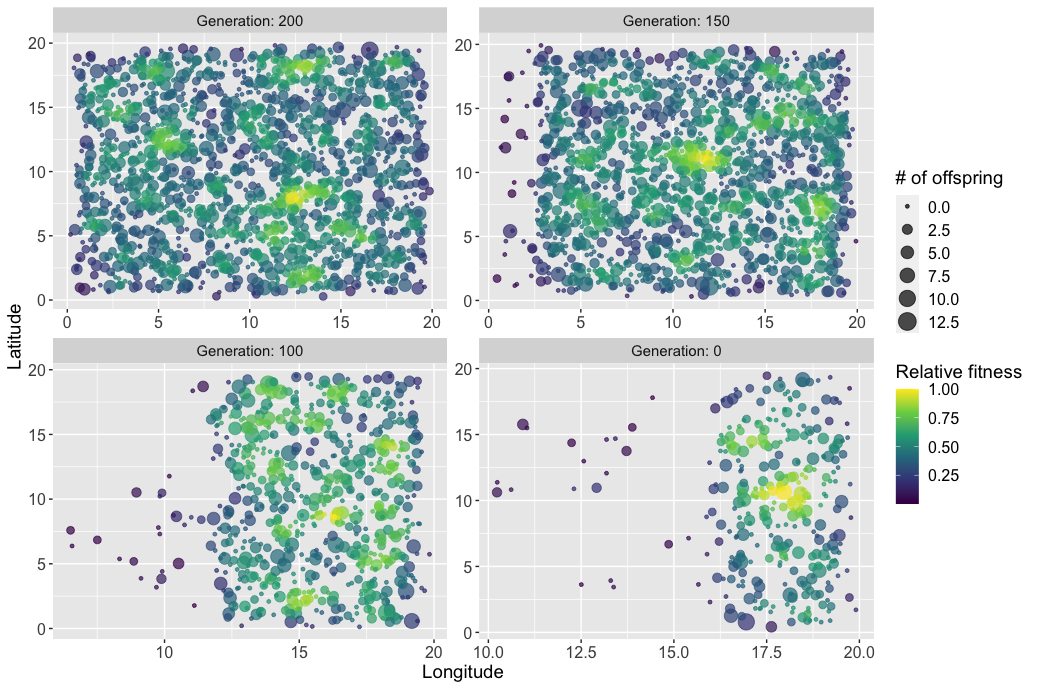

Figure S7. Spatial patterns of π in the *fragmentation* model before the contraction and 25, 50, and 100 generations after. Relative π is scaled to the pre-contraction levels, with values ~0.5 representing the mean. Size of the circle is scaled to absolute π.

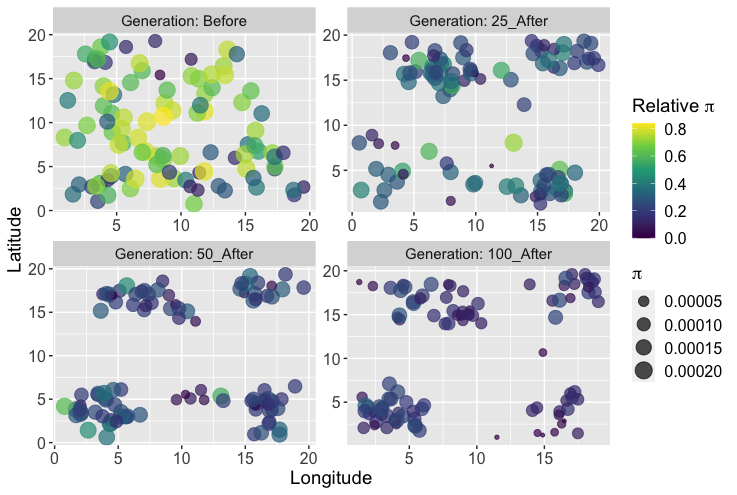

Figure S8. Shifts in relative fitness in the *fragmentation* models at generations 200, 150, 100, and 0.

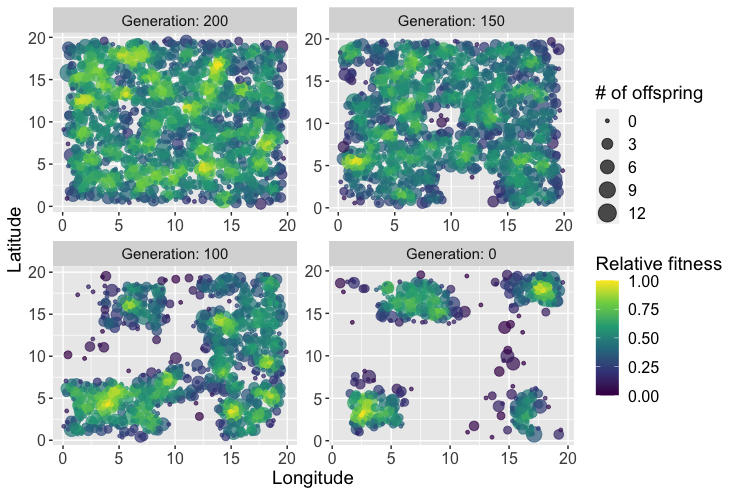

Figure S9. Spatial patterns of π in the *hollow* model before the contraction and 25, 50, and 100 generations after. Relative π is scaled to the pre-contraction levels, with values ~0.5 representing the mean. Size of the circle is scaled to absolute π.

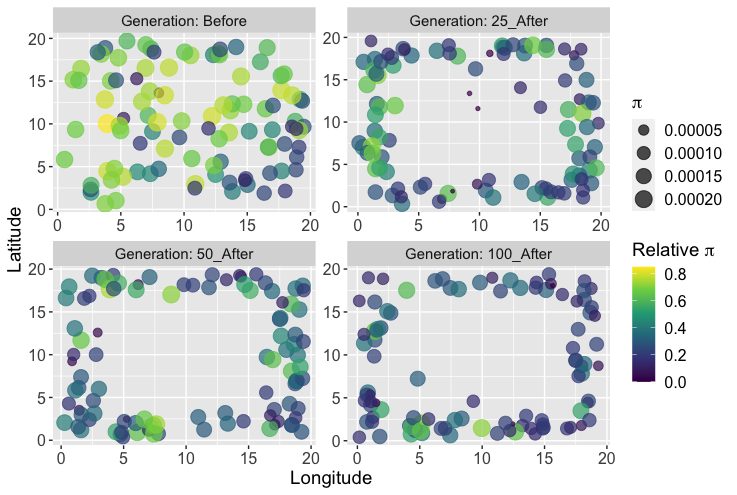

Figure S10. Shifts in relative fitness in the *hollow* models at generations 200, 150, 100, and 0.

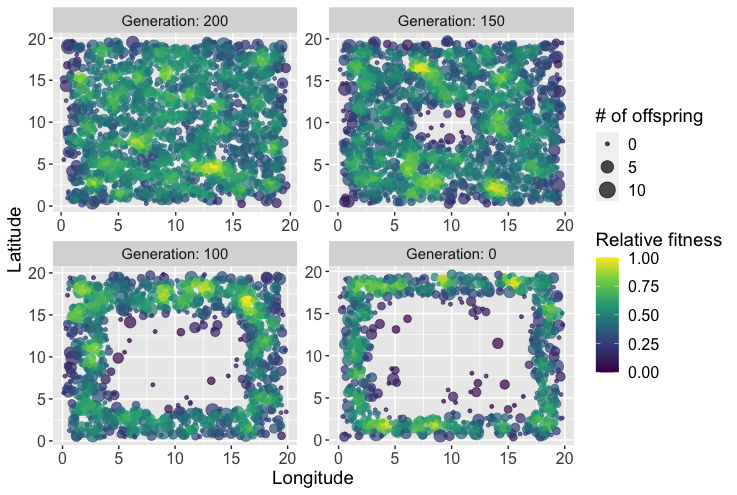

Figure S11. Sampling strategies from Tables S2–S13. Colors represent sampled groups for the *shrinkage* model.

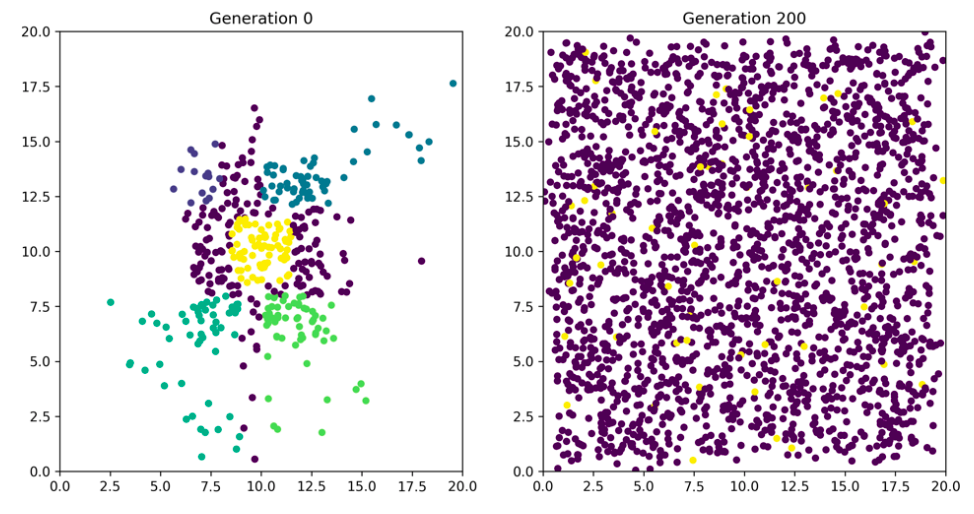

Figure S12. Sampling strategies from Tables S2–S13. Colors represent sampled groups for the *amputation* model.

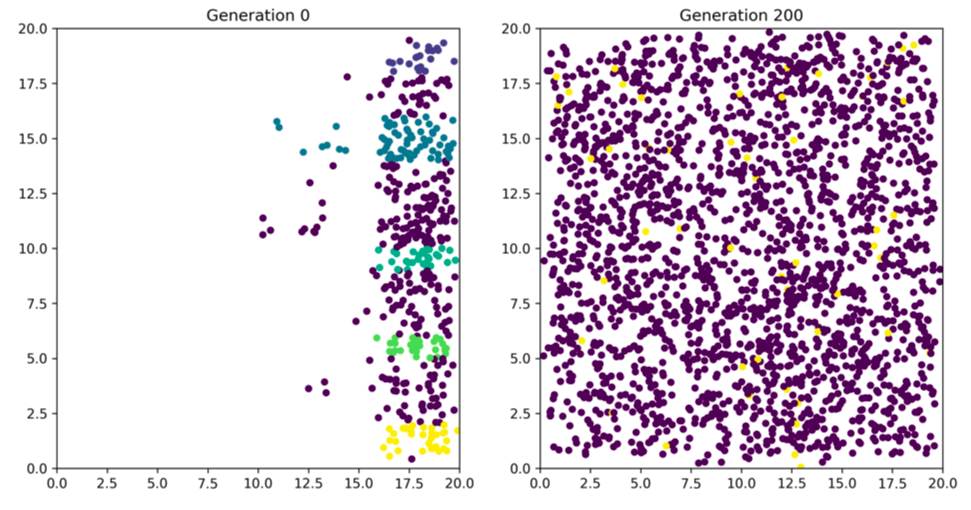

Figure S13. Sampling strategies from Tables S2–S13. Colors represent sampled groups for the *fragmentation* model.

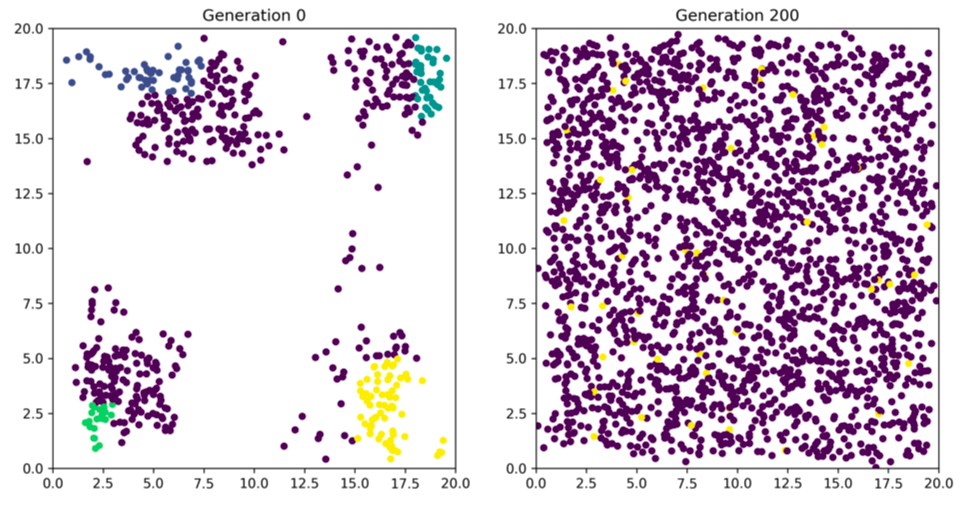

Figure S14. Sampling strategies from Tables S2–S13. Colors represent sampled groups for the *hollow* model.

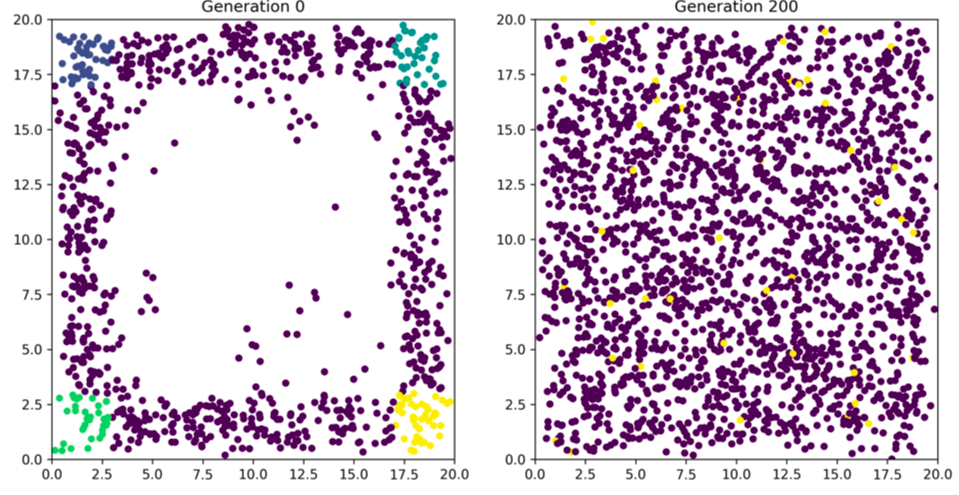
